# Environmental DNA as a complementary tool for biodiversity monitoring: A multi-technique and multi-trophic approach to investigate cetacean distribution and feeding ecology

**DOI:** 10.1101/2024.03.11.584480

**Authors:** L. Afonso, J. Costa, A. M. Correia, R. Valente, E. Lopes, M. P. Tomasino, Á. Gil, C. Oliveira-Rodrigues, I. Sousa-Pinto, A. López, C. Magalhães

## Abstract

The use of environmental DNA (eDNA) to assess the presence of biological communities has emerged as a promising monitoring tool in the marine conservation landscape. Moreover, advances in Next-Generation Sequencing techniques, such as DNA metabarcoding, enable multi-species detection in mixed samples, allowing the study of complex ecosystems such as oceanic ones. We aimed at using these molecular-based techniques to characterise cetacean communities, as well as potential prey in the northern coast of Mainland Portugal. During seasonal campaigns, we collected seawater samples, along with visual records of cetacean occurrence. The eDNA extracted from 64 environmental samples was sequenced in an Illumina platform, with universal primers targeting marine vertebrates. Five cetacean species were identified by molecular detection: common dolphin (*Delphinus delphis*), bottlenose dolphin (*Tursiops truncatus*), Risso’s dolphin (*Grampus griseus*), harbour porpoise (*Phocoena phocoena*) and fin whale (*Balaenoptera physalus*). Overall, except for the fin whale (not sighted during the campaigns), this cetacean community composition was similar to that obtained through visual monitoring, and the complementary results suggest their presence in the region all year round. In addition, the positive molecular detections of *B. physalus* are of special relevance since there are no visual records reported in the area. The detection of multiple known preys of the identified dolphins indicates they use these coastal areas for feeding purposes. While this methodological approach remains in a development stage, the present work highlights the benefits of using eDNA to study marine communities, with specific applications for research on cetacean distribution and feeding ecology, ultimately serving as the baseline of a methodological approach for biodiversity monitoring and marine conservation.

## Introduction

Environmental DNA (eDNA) is an emerging tool for biodiversity monitoring that has been gaining prominence in scientific research during the second half of the 21st century, with increasing numbers of scientific outputs being published every year [1]. For marine ecosystems, the application of eDNA detection methodologies is still in its infancy, although it has steadily evolved in the last decades given the interest in its potential [2]. Among the multiple applications, these allow verifying environmental health by studying microbial communities, characterising and quantifying stocks of marine vertebrates, such as teleost fish which represents vital knowledge for good management of the fisheries sector, and determining the presence and abundance of elusive species, such as cetaceans, allowing a greater effectiveness of the monitoring work [3–5]. Additionally, technological advances in Next-Generation Sequencing (NGS) techniques, via DNA metabarcoding, allow for simultaneous multi-species detections in environmental samples, permitting the study of multiple trophic levels within the same samples [6–8].

Cetaceans are widely dispersed mammals that inhabit most marine environments, from coastal habitats to neritic waters [9], playing a key ecological role in maintaining the balance of these ecosystems [10]. As keystone species, the conservation of cetaceans is often a top priority in international agreements, especially considering the anthropogenic threats they are currently facing [11]. Addressing the impacting issues in cetacean ecology is crucial, however, obtaining data that provides a detailed understanding of these animals is rather complex. Cetaceans are elusive individuals, spending the vast majority of their time underwater. Also, their distribution range is often very extensive, including areas where access for sampling is difficult due to logistical and financial limitations inherent to the marine wildlife monitoring work or even legal constraints [10]. Therefore, the development of new non-invasive methodologies, such as the metabarcoding analysis of eDNA samples, especially in complement to visual monitoring is a promising advance in the optimisation of monitoring effectiveness towards the better understanding of these highly complex species [10].

Especially regarding the use of eDNA samples for cetacean monitoring, few published studies were focused on marine mammals (see review on eDNA application to cetacean monitoring under [12]). Nevertheless, there are already successful case studies where it has been possible to identify a variety of cetacean species through environmental samples using both species-specific [5;13-16] and universal primers [4;17-23]. The possibility of detecting multiple species within the same environmental samples, enabled through metabarcoding, allows for a multi-trophic analysis that widens the utility of the samples for the monitoring of several taxa and application to various fields of research, including the study of trophic chains and species feeding ecology [23,24].

In the present study, a molecular detection methodology was developed, using eDNA samples as a tool for biodiversity monitoring, especially applied to cetacean species in a coastal region of the North of Portugal, located in the Eastern North Atlantic (ENA). The ENA region is an area of great interest regarding the diversity and abundance of cetaceans, with several different species of dolphins and whales being recurrently reported over the years [25–34]. Specifically, on the northwest Iberia, there is a high richness of cetacean species that feed on living marine resources [26;35;36]. Here, we sought to obtain additional and concrete data on the occurrence of cetaceans in this area and infer the ecological reasoning behind it by recurring to a universal approach to perform a multi-trophic analysis. Furthermore, we compared the eDNA results on cetacean species detection with the data obtained by traditional visual monitoring techniques, in order to assess the true potential of eDNA as a complementary tool across the panoply of methods employed for cetacean monitoring.

## Methods

### Study Area

Surveys to collect eDNA samples were performed in the north coast of Continental Portugal. This subregion, located in the northwest of the Iberia Peninsula, is of particular ecological interest due to the upwelling phenomenon strongly present along the coastline, thus enhancing primary production and providing great conditions for the development of complex and rich trophic chains [37–39]. In addition, the area is part of a particularly dynamic coastal region with several estuaries of rivers that flow into it [40]. Topographically, the study area is entirely placed on the continental shelf, with a relevant structure in the vicinity offshore, the Porto canyon [41] (Fig. 1).

**Fig. 1.**
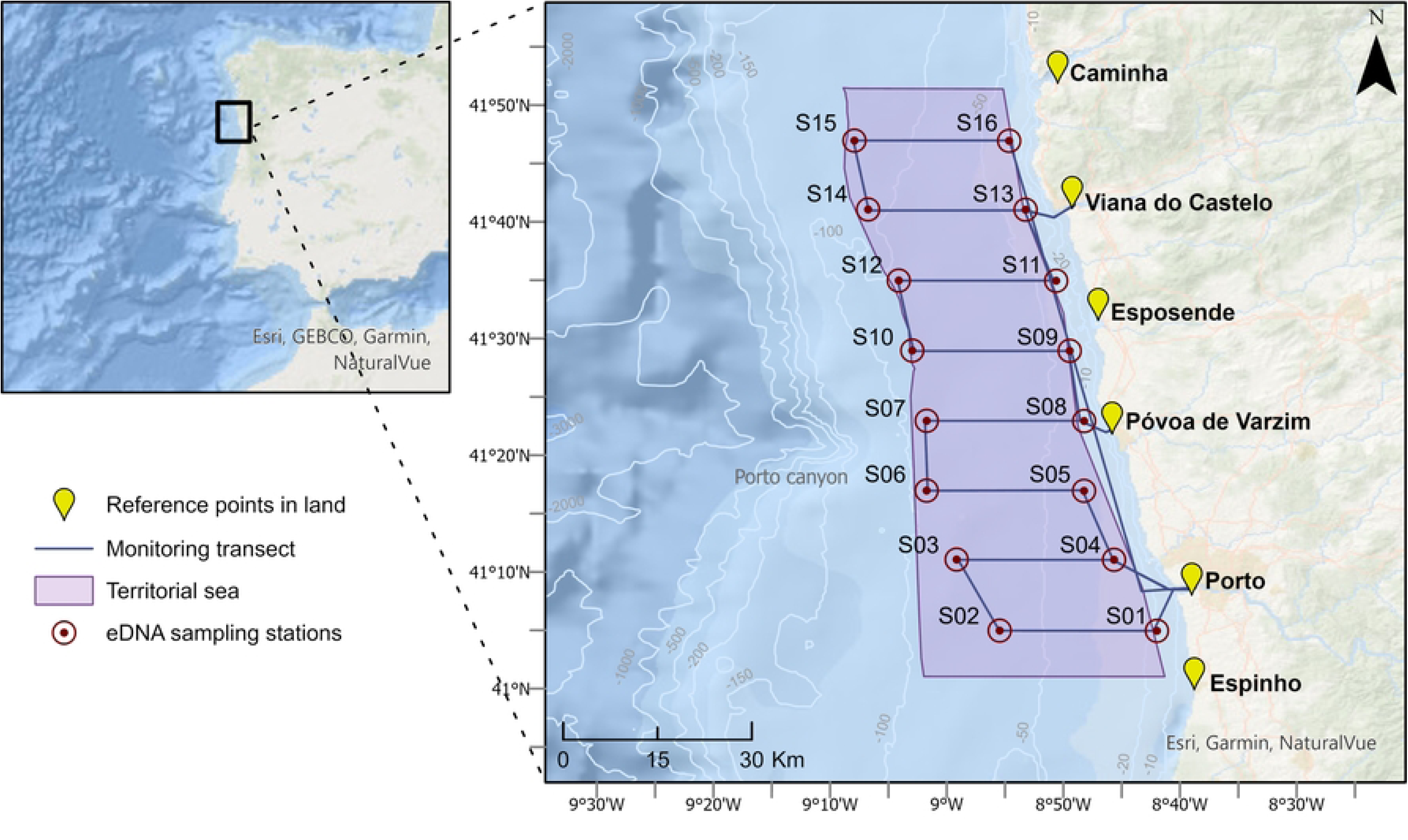
– Study area with surveyed transect of at-sea monitoring campaigns, conducted between the summer of 2021 and the winter of 2022/2023, in the north coast of Continental Portugal. Isobaths with bathymetry in meters.

In order to survey the area, four seasonal monitoring campaigns were carried out between the summer of 2021 and the winter of 2022/2023. Each campaign consisted of a survey transect with eight equidistant parallels, perpendicular to the coastline, spaced by approximately 6 nm, and covering distances of about 10 nm (approximately between 2 to 12 nm from the coastline), as shown in Fig. 1. For the collection of visual monitoring data, a previously established protocol [32] was followed to sample the occurrence of cetacean species sighted along the established transect (Fig. 1). The transect was designed so as to have an equitable range of observation capacity at all its points, guaranteeing a correct and complete visual monitoring of the study area. The collected visual monitoring data was imported into ArcGIS Pro for spatial analysis.

### Water Collection and Filtration

The environmental samples were sampled at pre-defined stations at the vertices of the campaign transect (Fig. 1, Table S1), using a bucket and a rope to collect 5 litres of water. Before water sampling, the samples were poured into 5L-containers. All the materials used for sample collection were previously cleaned with 10% bleach, rinsed with MilliQ water, and washed with local seawater (i.e., seawater at the sampling station) just before sample collection. After collection, volumes ranging from 1 to 3 litres were filtered immediately on board through Sterivex units (0.22 μm) using a peristaltic pump. After filtration, the samples were kept in liquid nitrogen, and stored at -80°C upon arrival at the laboratory. In total, 64 samples were collected, 4 at each station (two summers and two winters), with few exceptions: (1) two additional collections were made at S09 and S10 during the first summer campaign (2021); (2) during the first winter campaign, it was not possible to collect samples at stations S7 and S8, due to adverse weather conditions (Table S2).

### DNA Extraction

Total eDNA was isolated using the DNeasy® PowerWater® Sterivex™ Kit (QIAGEN), following the manufacturer’s instructions, with some adjustments to increase the DNA yield, namely: increased vortex times (10 minutes at all vortex steps) and 5 minutes rest time before the last centrifugation step. After extraction, DNA concentration for all samples was quantified using Qubit™ dsDNA High Sensitivity (HS) assay kit (Invitrogen™). Environmental DNA extraction was performed in a specifically dedicated laboratory for the extraction of genetic material, with the bench being always cleaned with ethanol prior to its use. All materials used were also sterilised in UV light before the start of the extraction process.

### Library Preparation, Sequencing and Bioinformatic Analysis

All samples were sequenced in high throughput sequencing in an Illumina MiSeq300 platform, using MarVer3(A) primers [8] and the Superfi II Polymerase (Invitrogen™). For marine vertebrate library preparation, a fragment of the vertebrate mitochondrial 16S rRNA gene was amplified and reamplified. In the first amplification step, PCRs were carried out in triplicate in a final volume of 10 μL, containing 2 μL of template DNA, 0,5 μM of the primers, 1X Platinum SuperFi II DNA Polymerase (Invitrogen), 0,8 mM dNTPs, 1X SuperFi II Buffer, 1X CES [42], and ultrapure water up to 10 μL. The PCR protocol was the following: an initial denaturation step at 98°C for 30 seconds, followed by 35 cycles of 98°C for 10 seconds, 49 °C for 10 seconds, 72°C for 30 seconds, and a final extension step at 72 °C for 5 minutes. Triplicate PCR products were pooled together. The oligonucleotide indices that are required for multiplexing different libraries in the same sequencing pool were attached in a second amplification step with identical conditions but only 5 cycles and with an annealing temperature of 60°C. Libraries were then purified using the Mag-Bind RXNPure Plus magnetic beads (Omega Bio-tek), following the instructions provided by the manufacturer. Lastly, the pool was sequenced in a fraction (3/8) of a MiSeq PE300 flow cell (Illumina). DNA metabarcoding library preparation and sequencing were carried out by AllGenetics & Biology SL (www.allgenetics.eu).

Following that, sequencing adapters were removed using Cutadapt v3.5 (Martin, 2011) and the originated reads went through a DADA2 [43] tool pipeline to remove other non-biological DNA sequences (e.g., primers), filter the reads according to their quality, denoise, dereplicate, cluster the resulting sequences into Amplicon Sequence Variants (ASVs), merge corresponding forward and reverse reads, and remove chimeric sequences. The taxonomic assignment of each ASV (with a minimum of 5 reads) was performed by querying its representative sequence against a local instance of the NCBI’s Nucleotide database (last updated on 18/09/2023), using the algorithm BLASTn v2.13.0+ [44] with the following parameters: percent identity of 99%, e-value of 1e-05, and a minimum hit coverage of 80%. If, based on these parameters, a matched ASV represented multiple species, the one with 100% identity was considered. For this part of the work, the ASVs corresponding to all cetacean species and Actinopteri superclass species with known occurrence in Continental Portugal (as of the Ocean Biodiversity Information System [OBIS] up to 05/09/2023) were selected. For the construction of the databases, all available nucleotide sequences (containing the 16S rRNA gene) from selected taxa were retrieved from the NCBI nucleotide database. Then, using the Geneious® software (v.7.0.6), primer sequences (forward and reverse) were annotated (using the ‘Add Primers to the Sequence’ tool), sequences were cut (using the ‘Extract PCR product’ tool) so they only contain the fragments spanning (and including) the PCR primers, and all nucleotide sequences not containing the entire fragment of interest were excluded from the database. Positively matched ASVs of each sample were then grouped into the respective sampling stations and seasonal campaigns for clear visual representation of the results. Representative stacked bar plots of identified taxa relative abundance and Actinopteri ASVs heatmaps for winter and summer monitoring campaigns were produced using the *ggplot2* (v.3.4.3) package [45], in R.

### Sampling Statistical Tests

In order to better understand whether there are significant variations in the concentrations obtained from samples taken in the different seasonal campaigns, Kruskal-Wallis chi-squared tests were carried out, and if significant differences were found (p-value < 0,05), we followed with a pairwise Wilcoxon test. The Wilcoxon test was also applied to compare the concentrations obtained from stations located at different distances from the coast (2 nm versus 12 nm). For this comparative analysis of DNA quantification results, samples that had DNA concentration below the Qubit’s HS kit detection limit (0,005 ng/µL) were assigned a value of 0,001 ng/µL. A sample log of all collected and extracted samples can be found in Table S2.

Statistical tests (Kruskall-Wallis test and pairwise Wilcoxon tests, with the level of significance set to 0,05) were carried out to understand the variation in the total number of ASVs obtained in the samples at the different seasonal campaigns and at the different distance from the coast (2 nm versus 12 nm).

## Results

### Visual Surveys

A total of 71 sightings of 4 different cetacean species were recorded during monitoring campaigns (Fig. S1A, Table S3). The majority of these records were from common dolphin (*Delphinus delphis*) with 45 sightings across the entire surveyed area, representing approximately 63.4% of all records. As for the other 3 species - bottlenose dolphin (*Tursiops truncatus*), Risso’s dolphin (*Grampus griseus*) and harbour porpoise *(Phocoena phocoena*) – these were sighted on 3 different occasions each. Besides the identified records to the species level, there were also 13 sightings of unidentified Delphinidae (18,3% of total records) and, on 4 occasions, it was only possible to identify cetacean occurrence at the superorder level (Cetacea). All sightings recorded with the respective date and geographical coordinates can be found in Table S3.

### eDNA Sampling

Among the campaigns conducted, the summer of 2021 showed much higher DNA concentrations compared to the other seasonal campaigns, with an average quantification of 9.46 ± 9.05 ng/µL (Median= 9,22 ng/µL). This was followed by the winter of 2021/22 with 5.10 ± 6.11 (Median= 3.02 ng/µL), the winter of 2022/23 with 2.26 ± 3.50 ng/µL (Median= 0.8 ng/µL) and, finally, the summer of 2022 with 1.08 ± 2.38 ng/µL (Median= 0.67 ng/µL). Statistical tests showed significant differences between the quantified concentrations of the different campaigns (p-value = 9.63^e-06^), these being explained by differences between summer 2021 and summer 2022, summer 2021 and winter 2022/23, and also between summer 2022 and winter 2022/23 (Table S4A). At the seasonal level, the summer shows higher concentrations, with 5.52 ± 8.18 ng/µL (Median= 3.51 ng/µL) compared to the 3.59 ± 5.02 ng/µL (Median= 1.49 ng/µL) obtained for the winter. However, this difference was not statistically significant (p-value = 0.79). Taking distance into account, the sampling stations near the coast (2 nm) had an average DNA concentration of 5.48 ± 8.18 ng/µL (Median= 3.2 ng/µL), while the DNA obtained in stations at 12 nm was quantified as 3.75 ± 5.21 ng/µL (Median= 1.26 ng/µL) on average. These differences were not statistically significant (p-value = 0.45).

### eDNA Monitoring: Sequencing Output and Taxonomic Identification

A total of 5 876 226 reads were generated by the Illumina platform with an approximate average of 90 403.48 ± 42 311.20. After quality-filtering steps, the final output was 3 735 272 total reads with an average of approximately 57 465.72 ± 40 520.33 (representing 65.6% of the input). There were 5 outlier samples that produced less than 50 reads. A total of 173 156 representative ASVs (of previously determined marine vertebrate species: cetaceans and selected Actinopteri species with known occurrence in Continental Portugal). Among the different seasonal campaigns, winter 2022/23 obtained the highest number of resulting ASVs with an average of 7 951.5 ± 23 991.69 (Median = 39), followed by winter 2022/23 with 2024 ± 2 980.45 ASVs (Median = 722.5), the summer 2021 with 932.33 ± 1 716.23 ASVs (Median = 201.5), and finally the summer 2022 with an average of 51.5 ± 178.56 ASVs (Median = 0). The statistical tests carried out indicate that there was a significant difference in the uptake of target eDNA taxa, both between seasons (p-value = 0.019) and between the different campaigns (p-value = 0.009). Pairwise tests highlight differences between the two summer campaigns, and both winter campaigns with the summer of 2022 (Table S4B). With regard to distance from the shore, samples collected closer (2 nm) resulted in an average number of ASVs of 988.13 ± 2087.06 (Median = 0), while those further away (12 nm) resulted in 4,423.31 ± 17,126.03 ASVs (Median = 52.5). However, differences of obtained ASVs obtained in relation to the distance to coast were not significant (p-value = 0.64).

Of the 64 eDNA samples sequenced through metabarcoding, 33 (51.6%) had positive detections of representative ASVs, from which 10 of the samples (15.6%) had positive detections of cetacean ASVs. By grouping the results by sampling station, we noticed that all sampling stations had positive detections for either Cetacea or Actinopteri. Relative abundances of all cetacean species identified, and Actinopteri taxa detected in 4 or more stations, are illustrated in Fig. 2.

**Fig. 2.**
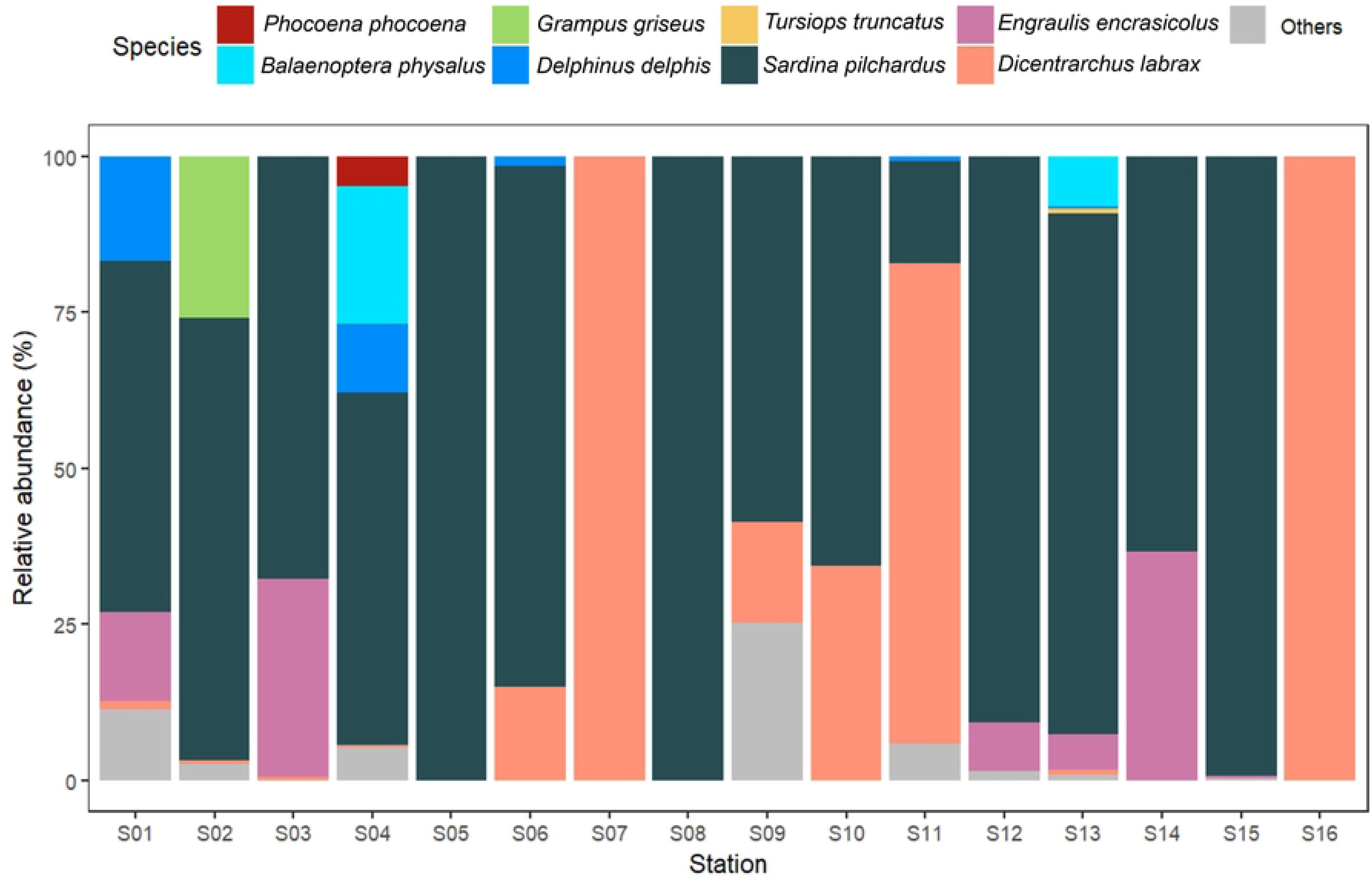
- Taxonomic assignment from environmental DNA samples collected in 16 sampling stations, in the North coast of Mainland Portugal, between 2021 and 2023. The plot represents the relative abundance (%) of sequences matched for each marine vertebrate species, considering pre-determined Cetacea, and Actinopteri species with positive detection in 4 or more stations (see methods). “Others” represents Actinopteri superclass species identified in 3 or fewer stations.

Focusing on cetaceans, 5 different species were detected, with at least one of them identified in 7 out of the 16 stations (Fig. 2 and Table S4). The common dolphin was the most detected species, being identified at 6/16 stations, followed by the harbour porpoise and the fin whale (*Balaenoptera physalus*), occurring at 2/16 sampling stations. The Risso’s dolphin and the bottlenose dolphin were detected on 1/16 stations.

Regarding Actinopteri species, 19 species were identified, with at least one of them detected in all sampling stations (Fig. 2 and 3). Of all these detections, the most notable result was the highly frequent presence of the sardine *(Sardina pilchardus*) - in 14/16 sampling stations; the European seabass (*Dicentrarchus labrax*) - in 9/16 stations; and the European anchovy (*Engraulis encrasicolus*) - in 6/16 of the sampled sites (Fig. 2 and 3). All the Actinopteri species identified, as well as the respective number of ASVs detected at each station, can be found in Table S4. Additionally to the frequency of the detections across sampling stations, there is also a higher abundance of captured ASVs of the species *S. pilchardus* and *D. labrax* (Fig. 3). In 5/16 sampling stations, 100% of the matched ASVs belonged to either *S. pilchardus* or *D. labrax* (Fig. 2). The highest number of ASVs was obtained for *S. pilchardus* in Station S15 (off north of Viana do Castelo) (Fig. 5). Station S01 (southwest of Porto, at ∼2nm from coast) had positive detections for a higher number of selected Actinopteri species in both seasons (Fig. 3). The Station S08 (near Póvoa de Varzim) only had one minor detection (9 BLAST hits for its respective ASV, see Table S5) for *S. pilchardus* across all monitoring campaigns.

**Fig. 3.**
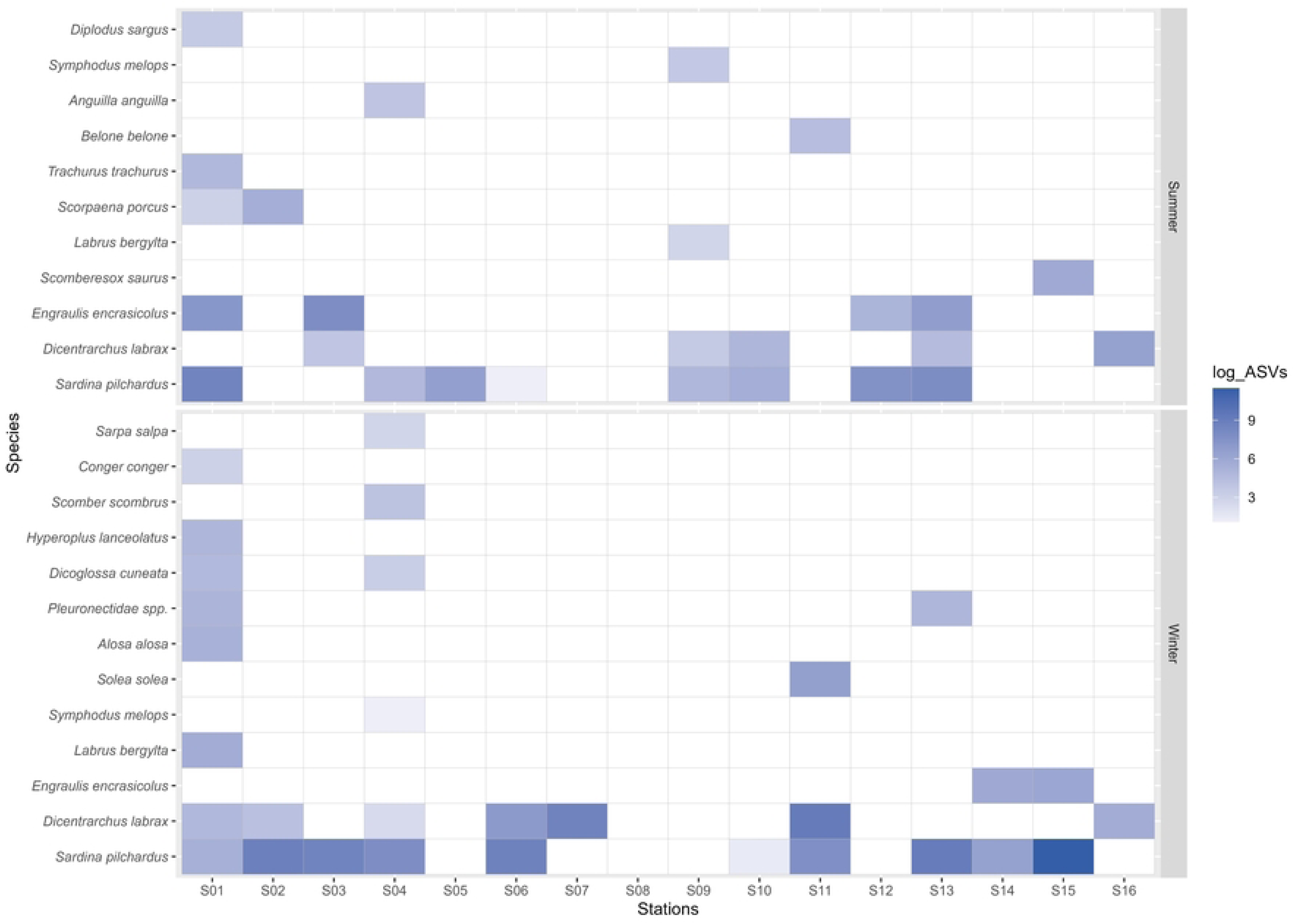
- Number of Amplicon Sequence Variants (ASVs) detected for each species of the Actinopteri superclass molecularly identified in environmental DNA samples, collected at the sampling stations surveyed in the summer (top) and winter (bottom) in the North coast of Mainland Portugal, between 2021 and 2023. The number of ASVs was logarithmised for scaling purposes.

### Cross-checking eDNA with Visual Monitoring Data

Overall, by cross-referencing the data obtained by the two sampling methodologies applied in this study, we can verify the occurrence of the following four species in both datasets (visual records, eDNA molecular detection): *D. delphis*, *P. phocoena*, *T. truncatus* and *G. griseus*. In addition to these, positive molecular detections of *B. physalus* were obtained through eDNA analysis. Regarding the spatial distribution of these records, it is clear that *D. delphis* was sighted not only in greater abundance but also with a dispersed distribution throughout the region studied, which is also in line with the data obtained by molecular detection. For the other identified species, their DNA abundance was lower and there were fewer visual records obtained as well.

Overall, no pattern of spatial or temporal correlation between molecular detections and visual records is clear. An inter-seasonal comparison of the spatial distribution of the records obtained by the visual surveys with the eDNA detections reveals contrasting results (Fig. 4). In terms of composition of the cetacean community, in the summer campaigns, all the 4 species mentioned above were sighted; while in the winter campaigns, sightings of *D. delphis* accounted for all records, excepting one sighting of *T. truncatus* (considering sightings identified at the species level) (Fig. 4A). Through eDNA analysis though, all the species were molecularly detected in the winter campaigns (including also *B. physalus*), while in the summer season the only species identification through molecular detection of cetacean species was of *D. delphis* (Fig. 4B). Disregarding seasonality and accounting for spatial distribution only, the location of molecular detections of each species at the sampling stations seems to align better with the visual records at the latitudinal level than in relation to distance to the coastline (i.e., longitudinally). In summary, it is worth highlighting that: i) in both datasets, *D. delphis* was found to be distributed across the entire area with higher frequency north of Póvoa de Varzim; ii) *T. truncatus* was distributed in the north of the study area, visually recorded at different distances from the coast but only detected with molecular methods in the most coastal sampling station (S13); iii) *P. phocoena* was detected in both datasets south of Póvoa de Varzim, and mostly in stations near coast (with only one sighting further from the coast); iv) the molecular detections of *B. physalus* occurred in the stations closer to the coast, one in the north and another in the south of the study area (Fig. 4B).

**Fig. 4.**
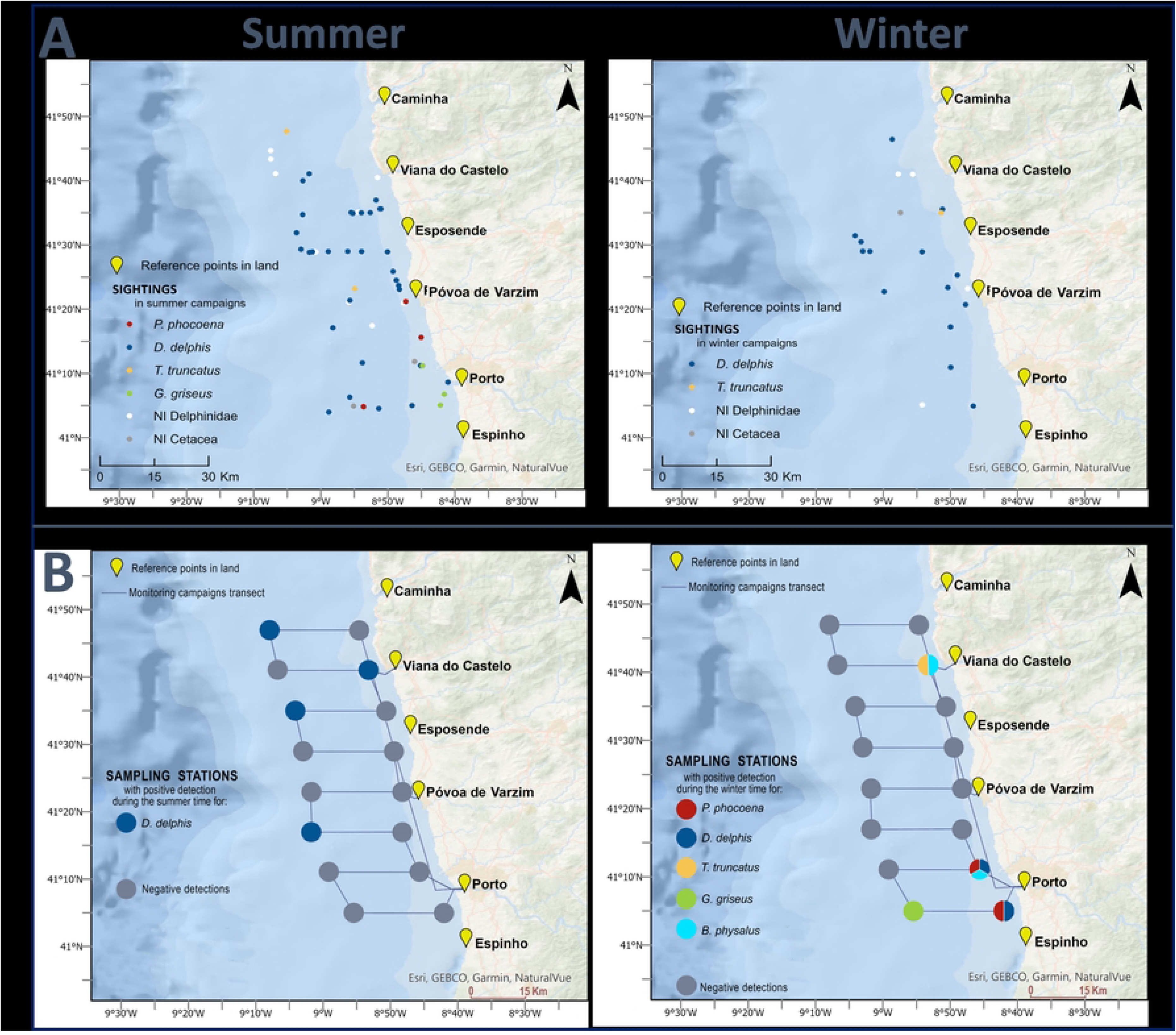
– Seasonality of cetacean records obtained by visual monitoring (A) and molecular analysis of environmental DNA (B), for summer (left) and winter (right) campaigns carried out in the North coast of Mainland Portugal, between 2021 and 2023.

### Multi-trophic Analysis

Across all campaigns, multiple Actinopteri detections coincided with the molecular presence of cetacean species (Table 1). *D. delphis* was detected by eDNA simultaneously to *S. pilchardus* (on two occasions), *B. belone*, *E. encrasicolus*, *D. labrax*, *S. saurus* and *L. bergylta*. *P. phocoena* detections were coincident with a higher richness of Actinopteri species, including *S. pilchardus*, *D. labrax*, *S. scombrus*, *L. bergylta*, *Pleuronectidae spp.*, *H. lanceolatus* and *C. conger* (Table 1). *G. griseus* and *B. physalus* were also detected at the same time as *S. pilchardus* (Table 1), while *T. truncatus* was solely detected without associated potential Actinopteri prey.

**Table 1.** Coincident Cetacea and Actinopteri species detections by environmental DNA at sampling stations surveyed across seasonal monitoring campaigns carried in the North coast of Mainland Portugal, between 2021 and 2023.

| Monitoring campaign | Sampling station | Cetacean | Actinopteri |
| --- | --- | --- | --- |
| Summer 2021 | S11 | *Delphinus delphis* | *Belone belone* |
|  | S13 | *Delphinus delphis* | *Sardina pilchardus*<br>*Engraulis encrasicolus*<br>*Dicentrarchus labrax* |
|  | S15 | *Delphinus delphis* | *Scomberesox saurus* |
| Winter 2021/22 | S01 | *Phocoena phocoena* | *Sardina pilchardus*<br>*Dicentrarchus labrax*<br>*Labrus bergylta*<br>*Alosa alosa*<br>*Pleuronectidae spp.*<br>*Dicoglossa cuneata*<br>*Hyperoplus lanceolatus*<br>*Conger conger* |
|  | S04 | *Phocoena phocoena* | *Sardina pilchardus*<br>*Dicentrarchus labrax*<br>*Alosa alosa*<br>*Dicoglossa cuneata*<br>*Scomber scombrus*<br>*Sarpa salpa* |
| Winter 2022/23 | S01 | *Delphinus delphis* | *Labrus bergylta* |
|  | S02 | *Grampus griseus* | *Sardina pilchardus* |
|  | S04 | *Delphinus delphis*<br>*Balaenoptera physalus* | *Sardina pilchardus* |

## Discussion

### Environmental DNA Sampling and Uptake

The sampling techniques here employed were chosen due to their on-field practicability (collection of seawater and filtration) and lower risk of contamination (no direct handling of filters during DNA extraction). To date, there are no generalised standard protocols for the detection and identification of cetacean species through eDNA, with a wide variation of methods currently being implemented [12]. The lack of a standardised, reliable, and repeatable protocol for sample collection (and subsequent processing and analysis) still constitutes a major obstacle to the use of this monitoring tool for marine biodiversity monitoring [46].

On the northwest coast of the Iberian Peninsula, upwelling phenomena tends to occur in the summer, inducing an increase in primary productivity and subsequent development of the trophic chain [39]. Therefore, it was expected DNA concentrations to be uniformly higher in the samples collected during summer campaigns, which was not the case. However, while the 2021 summer campaign presented the highest values of DNA concentration, which could be explained by the mentioned oceanographic phenomena, the exact opposite was observed in 2022. This significant difference, although not ruling out possible human error in the different sampling steps, may be due to some unknown oceanographic event happening during sampling days or to other environmental variables with known impact on DNA preservation, such as temperature [47;48], pH [49;50] and/or UV radiation exposure [51]. The lack of inter-seasonal significant differences in the DNA concentration obtained from eDNA samples, in contrast to the significant inter-campaign differences observed, may evidence that the seasonal (i.e., inter-annual cycles such as seasonal upwelling) influence is less relevant than the impact of less cyclic and more ephemeral phenomena on the DNA preservation and retrieval. Regarding the number of target ASVs, significant differences found between both seasons and monitoring campaigns, suggest a higher target DNA uptake in winter months.

Nevertheless, this result should be interpreted with caution, since these differences are mainly justified by the low numbers evidenced in the summer 2022 campaign (for which a very low DNA concentration was obtained). As far as the proximity to the coast, although with some variation, there were no significant differences in terms of DNA concentrations or the number of ASVs obtained. Researchers have reported a faster DNA degradation in inshore areas associated with intense anthropogenic pressures [52]. Nevertheless, coastal areas are also likely to have higher biomass, in comparison with offshore areas, due to the coastal upwelling phenomena [39] and the input of the rivers [53;54]. The coastal waters of the north coast of Continental Portugal are heavily influenced by the input of various fluvial water masses such as the Douro, Cávado, and Minho rivers [55;56], which could also be interfering with sampling results. The river plume of the Douro river is known to vary substantially inter annually, with increasing extension into the ocean in winter months [57]. Overall, we consider that further studies are required in the future to monitor the influence of environmental conditions on eDNA uptake, and therefore evaluate sampling success in dynamic marine coastal ecosystems.

### Bioinformatic Analysis Criteria

To compile the BLASTn databases, we established a strict criterion, comprising the amplified fragment of interest and the sequences of the forward and reverse primers for the aforementioned species of interest. However, this type of approach for sequence selection has its pros and cons. On the positive side, it allows for greater certainty that a given sequence belongs to a respective species since smaller fragments can result in greater similarity between species (conservative approach). However, this method can lead to a loss of information by possibly removing sequences of interest. Moreover, and particularly for assembling the database of sequences corresponding to the Actinopteri species, the approach was only focused on the species already recorded along the Portuguese coast, substantially reducing the size of the database. Such a decision was made since, in this context, the purpose for the detection of Actinopteri species was tied to the ecological role they play in the diet of cetaceans.

Given the parameters applied in the BLASTn algorithm used for the taxonomic assignment (identity higher than 99%), there were only two cases in which multiple species matched to a given ASV: one within Cetacea taxa, and another within Actinopteri. A specific example of multi-species assignment to an ASV was a case where the ASV was assigned to *D. delphis* with 100% identity, but other taxa were assigned with a percentage identity of 99.14% (the genera *Tursiops* spp. and *Stenella* spp.). This was to be expected given the genetic similarity of these species pertaining to the oceanic dolphin family (Delphinidae), as described by McGowen et al. [58]. Especially between the common dolphin and the bottlenose dolphin, there are only two Single Nucleotide Polymorphisms (SNPs) on the amplicon obtained with the MarVer3 primers. Besides this case, there was a double identification with 100% percent identity for the same ASV - 7 sequences for the Atlantic horse mackerel (*Trachurus trachurus*) and 1 for the megrim (*Lepidorhombus whiffiagonis*) in the NCBI database. In this case, the species *T. trachurus* was selected as the correct identification. Such decision was because the matched sequence of *L. whiffiagonis* referred to samples collected in the Mediterranean Sea, in which case the authors of the work demonstrated a great variation in the mitochondrial 16S rRNA region between the Mediterranean and the Atlantic Ocean populations [59].

### Cetacean Detections: eDNA *versus* Visual Monitoring

As described in the results, the species detected through visual monitoring were also detected by eDNA, with the additional record of the species *B. physalus*, which was only recorded through molecular methods in the environmental samples. This is a species with non-existent sightings in this region of Continental Portugal, even though it has been recorded in other parts of the country’s coastline [60]. Additionally, in the Galician coastal waters, adjacent to the study area, the species occurs rather frequently [30;61;62]. Offshore Galicia, there are relevant topographical structures, seamounts, that may act as hotspots for pelagic biodiversity [63], such as the Galicia Bank [64]. The obtained detection of *B. physalus* DNA may be evidence of an occasional occurrence of the species in waters off northern Portugal.

As shown by the results, the common dolphin represented the vast majority of the sightings in the study area. The wide distribution and frequent presence of *D. delphis* along the entire northern Portuguese coast was corroborated by molecular detection on eDNA samples. The harbour porpoise was detected via eDNA at two stations located closer to the coastline. That result is in line with the frequent occurrence of the species at the mouth of the Douro River (located in proximity to the sampling sites), as described by Gil et al. [65] with occurrence records obtained through visual monitoring. As for *T. truncatus* and *G. griseus*, both species were only sighted on very few occasions. By eDNA, each of these was detected at one station, emphasising the less frequent occurrence in the north of Continental Portugal, in comparison with *D. delphis* and *P. phocoena*. *T. truncatus* was detected via eDNA in northern latitudes, where the sightings of the species were also registered. As for *G. griseus,* the DNA was detected in the southern quadrant of the study area, where the species was also recorded through visual monitoring, where they were also recorded by molecular detection. On the other hand, The bottlenose dolphin is a frequent species in the northwest of the Iberian Peninsula [66], therefore detection through visual and molecular methods was anticipated, although more records were to be expected. Overall, the combination of the results from visual monitoring and eDNA suggests the year-round presence of the dolphin and porpoise species on the north coast of Continental Portugal. Given the difficult weather conditions for visual monitoring in the ENA for a large part of the year, mostly in winter months, and consequently low monitoring effort in the region, complementary results from eDNA prove to be very useful to increase baseline knowledge of these coastal populations.

Besides multi-species detection on eDNA samples, the application of the methodology for single-species detection may be of particular importance. This is the case given the need to monitor resident populations in decline, as shown by Ma et al. [14] where eDNA tools were applied for conservation efforts. In the north coast of Continental Portugal, we believe that this approach is extremely pertinent for species of conservation interest, such as the harbour porpoise, whose population in the mainland Portuguese coastline is predicted to be extinct in 20 years (Critically Endangered, [67]).

Despite the aforementioned promising results, some challenges remain to be addressed for the future consistent application of this novel monitoring tool. In previous eDNA metabarcoding studies, it was usually not possible to distinguish between species of the same genus, as observed in Valsecchi et al. [22] for the *Tursiops* and *Stenella* genera, and even within the Delphinidae family [4;20]. In the present work, this was also a challenge during the assignment of ASVs, even though a conservative selection criterion was established. Therefore, this limitation represents a significant obstacle to meet monitoring objectives. In addition, although the use of universal primers may provide us with relevant information at the multi-trophic level, there is a major drawback when the technique is used to detect a certain taxon (e.g., monitoring cetacean occurrence). Besides the likelihood that primers bind to non-target DNA in greater abundance at the time of amplification, which can result in false negative detections [68;69], it is impossible to pre-select the samples with the target DNA before the sequencing step as the detections in the post-PCR electrophoresis may allude to different taxonomic groups. Thus, in the context of the applicability of eDNA for more efficient and accessible (i.e., cost-effective) taxon-specific monitoring, it is advisable to use taxon-specific primers. In the specific case of cetaceans, where, in addition, relative abundance may often be rather low (in comparison to other marine taxa), the development of cetacean-specific primers is one of the next key steps to address in the continued emergence of the application of eDNA tools for cetacean monitoring.

### Monitoring Multiple Trophic Levels

The introduction of universal primers in the study of marine communities allows the collection of data from multiple trophic levels, with posterior inference of possible ecological interactions and prey/predator relationships. Universal primers also provide a deeper understanding of unusual distribution patterns, as was the case in Zhang et al. [24] where the authors used data obtained by eDNA metabarcoding to assess the food resources of Eden’s whale (*Balaenoptera edeni edeni*), and in Visser et al. [70] where the cephalopod community composition was studied to better understand Risso’s dolphins (*G.griseus*) and Cuvier’s beaked whales (*Ziphius cavirostris*) foraging behaviours.

In this study, it was possible to detect numerous species from the Actinopteri superclass. In addition to the sardine, highly abundant and dispersed in the study region, and reported as main target for the diet of *D. delphis* on the Iberian coast [71;72], other species detected have also been found in its stomach contents in the portuguese continental coast, such as the European anchovy (*Engraulis encrasicolus*), the sole (*Soleidae spp.*), the needlefish (*B. belone*), the scads (*Trachurus spp.*) and the Atlantic mackerel (*Scomber spp.*) [72]. Another species with abundant and dispersed detections in this study, the European seabass, was also found in the stomach contents of *D. delphis* in the English Channel [73]. This latter coastal fish species, of high economic interest, is of particular relevance from a cetacean conservation perspective since its feeding preferences overlap with those of the dolphins, with reported implications for their bycatch by promoting interactions with the fisheries sector [74]. The aforementioned sardine is also extremely relevant to this topic due to its commercial importance in Portugal, being a main target of purse seine fisheries in the country [75]. In Continental Portugal, interactions between various dolphin species with this fishing method are frequent and can result in the bycatch of these animals [76;77]. In the case of *P. phocoena*, many of the species identified simultaneously, such as *Scomber scombrus* and the ballan wrasse (*Labrus bergylta*), and others occurring in other stations, such as *T. trachurus*, are also known prey for these animals in the ENA [78;79], which suggests that the population sighted regularly near the mouth of the Douro River [65] uses this coastal area for feeding purposes. Although not detected in coincidence, a wide variety of fish taxa identified in the studied area were found in *T. truncatus* stomach contents in Iberia, namely *S. pilchardus*, European conger (*Conger conger*), *E. encrasicolus*, *Trachurus spp.* and *Soleidae spp.* [80].

Therefore, data obtained from metabarcoding techniques evidenced the ecological importance of the region as a feeding area for coastal cetaceans, also highlighting the prey species available to their populations, which may have relevant implications for management and conservation strategies. Environmental DNA is thus proving to be a promising tool for multitrophic assessments, not only to study biodiversity occurrence, but also to infer about ecological processes and investigate trophic relationships.

## Conclusion

In summary, we demonstrated the potential of metabarcoding methods applied to eDNA samples, for biodiversity assessments, with special relevance as a cetacean monitoring technique for the study of cetacean distribution and feeding ecology. However, there are still obstacles and difficulties to overcome. More studies are necessary to better understand this novel sampling method, as there are still knowledge gaps in the application of the method, from environmental sampling to the analysis of the sequencing results.

Positive detections of cetacean species in this work constitute important data, as it was possible to characterise the northern coast of Continental Portugal in terms of cetacean occurrence, not only reproducing similar results but also complementing the data obtained through visual monitoring. Here, eDNA monitoring allowed us to conclude that dolphin and porpoise species that are less often sighted during visual surveys, even with known occurrence in the study area, have a probable all-year-round presence in this region of the ENA. Additionally, this study also shows the potential of this molecular-based technique to collect data when the monitoring effort is hampered by the inherent conditions of the sampled region (e.g., weather conditions), especially for less frequent species (possibly, without existing visual records). Furthermore, by expanding the analysis to lower trophic levels, the detection of multiple species known as cetacean prey species evidenced their use of these coastal areas for feeding purposes.

In conclusion, this work has demonstrated the potential of an innovative monitoring methodology for studying complex marine biological communities, such as cetaceans, even allowing for a multi-trophic approach, essential for conservation efforts. Therefore, although the effectiveness of using eDNA as a tool in cetacean monitoring programmes remains under development, this work represents a step forward towards that goal.

## Acknowledgements

Ph.D. fellowships for authors RV (SFRH/BD/144786/2019) and AG (PD/BD/150603/2020) were granted by Fundação para a Ciência e Tecnologia (FCT, Portugal) under the auspices of Programa Operacional Regional Norte (PORN), supported by the European Social Fund (ESF) and Portuguese funds (MECTES). This work is a result of the project ATLANTIDA (ref. NORTE-01-0145-FEDER-000040) supported by the Norte Portugal Regional Operational Programme (NORTE2020), under the PORTUGAL 2020 Partnership Agreement and through the European Regional Development Fund (ERDF); and of the project EMPHATIC funded by Biodiversa+, the European Biodiversity Partnership, under the joint call 2022 – 2023 BiodivMon for research proposals, co-funded by the European Commission and with the following funding organisations: Fundación Biodiversidad (FB, Spain), Fundação para a Ciência e Tecnologia (FCT, Portugal), Agence Nationale de la Recherche (ANR, France), and the Ministry of Universities and Research (MUR, Italy).

